# Asynchronous Detection of Erroneous Behaviors in Human-Robot Interaction with EEG: A Comparative Analysis of Machine Learning Models

**DOI:** 10.1101/2023.09.30.560271

**Authors:** Zhezheng Ren, Xuzhe Xia, Yuzhi Tang, Bo Zhao, Chun Pang Wong, Dongsheng Xiao

## Abstract

We present a comparative analysis of two distinct machine-learning models designed to detect asynchronous errors during Human-Robot Interaction (HRI). The models under scrutiny are a customized ResNet model and an ensemble model, both trained and validated using EEG data. The ResNet model is a unique adaptation of the Residual Network, comprising a one-dimensional convolutional layer followed by batch normalization and ReLU activation. It also features a custom Residual Block structure and an adaptive average pooling layer, concluding with a fully connected linear layer for binary classification. The ensemble model, on the other hand, incorporates various machine learning models such as MLP, logistic regression, SVM, random forest, and XGBoost, unified in a pipeline with feature extraction and transformation steps. A critical component of our research is the innovative probability map, which maintains a granularity of 0.1 seconds. This map forecasts the likelihood of forthcoming one-second intervals being classified as either Error or Non-error. Our comparative analysis reveals significant variations in the performance of the two models, both of which exhibit promising results in detecting erroneous behaviors during HRI. We provide detailed validation results, including the accuracy, F1 score, and confusion matrix for each model. This study offers valuable insights into the potential of machine learning in enhancing HRI efficiency and accuracy, indicating promising directions for future research.

## INTRODUCTION

The integration of robotics into everyday life is no longer a fantastical dream but an evolving reality. With robots and artificial intelligence becoming increasingly ubiquitous, the interaction between humans and machines is a field that has garnered significant attention. Human-Robot Interaction (HRI) stands at the intersection of technology, neuroscience, and human psychology, aiming to foster seamless communication between humans and robots(Ayanoğlu & Duarte, 2019; Dautenhahn & Saunders, 2011; Filippini & Merla, 2023; Henschel, 2020; Jost et al., 2020; Okamura et al., 2010; Oliveira et al., 2021).

Historically, the concept of HRI has been romanticized in science fiction literature, such as Isaac Asimov’s “Three Laws of Robotics” and even in modern representations like Marvel Comics’ robot healthcare companion – Baymax(Bartneck, 2004). Today, the rapid advancements in artificial intelligence and robotics have turned this fiction into tangible applications. Robots are now sharing spaces with humans, collaborating to achieve common goals, and the essence of this interaction lies in understanding and decoding human emotions and intentions(Urakami & Seaborn, 2023; Y. Wang, 2021; Xue et al., 2020).

The challenge of facilitating more natural and intuitive interaction between humans and robots has led to the exploration of human brain activity as a potential source of intrinsic feedback(Kim et al., 2017). By retrieving information from electroencephalogram (EEG) data, insights into the human mindset and subjective satisfaction with the robot’s performance can be obtained(Chiang, 2023; Tonin et al., 2021). This, however, poses tremendous challenges such as the feasibility of recording and using EEG data under real-world conditions(Protzak & Gramann, 2018), and decoding the brain asynchronously(Lotte et al., 2018). Addressing these challenges is central to the ongoing research efforts in the field. The main goal of the present challenge is to develop innovative signal processing and machine learning approaches for the asynchronous detection of erroneous behaviors based on single-trial EEG analysis. This involves the training of a machine learning model to detect the onset of deliberately introduced errors and perform validation using cross-validation techniques and unlabeled test data.

This study is at the forefront of these efforts, providing a comparative analysis of two distinct machine-learning models designed to detect asynchronous errors during HRI using EEG data. We present a customized ResNet model and an ensemble model, both rigorously evaluated and validated. We further introduce an innovative probability map, offering granularity in forecasting the likelihood of erroneous behaviors. Our analysis uncovers significant variations in the performance of the two models, highlighting the potential of machine learning to enhance HRI efficiency and accuracy.

Our study is further validated by our first-place win in the offline competition. The contributions of this research can be summarized as: (1) Customized and comparative analysis of ResNet and ensemble models for error detection in HRI. (2) Introduction of an innovative probability map for granular forecasting. (3) Significant insights into the potential of machine learning in enhancing HRI, signaling promising directions for future research.

By successfully intertwining the fields of neuroscience, robotics, and machine learning, this study sets a precedent for future work in HRI, demonstrating the potential to translate theoretical concepts into practical applications that can enhance human life and collaboration with robots.

## METHODOLOGY

### Experiments

The experiment was conducted as part of the IntEr-HRI competition at IJCAI 2023 (https://ijcai-23.dfki-bremen.de/competitions/inter-hri/), aiming to record a dataset for the detection of event-related potentials (ERPs) in EEG data(Kueper et al., 2023). The study took place inside the lab of the SMT department (BB311) at the University of Duisburg-Essen and involved a total of eight young, healthy subjects.

The subjects performed a flexion-extension task while standing and wearing an active orthosis on their right arm. This orthosis, designed to move automatically upon the application of small forces, executed flexion or extension movements with the right arm. Errors were deliberately introduced during normal operations to study the corresponding ERPs. Variations were introduced for specific subjects to explore different error positions and feedback mechanisms. The experiment was based on two primary conditions: (1) Baseline Condition: Subjects executed 15 flexions and 15 extensions under standard conditions, with the orthosis moving between fully extended (−10 degrees) and fully flexed (−90 degrees) positions. (2) Multimodal Condition: The same movements were executed, but errors were randomly introduced 6 times around specific mean error positions. Subjects were instructed to squeeze an air-filled ball if they recognized the error. The experiment included the collection of EEG and EMG data. Errors were manifested through a change of movement direction for 250 ms, and subjects’ responses were captured through their interactions with the air-filled ball.

The Brain products LiveAmp64 system was employed for EEG measurements. This system is wireless and utilizes active electrodes. Recordings were performed both wirelessly on the computer and on hardware using an SD-card. The BrainVision Recorder software (V. 1.25.0001) was used for the recording process. A total of 67 channels were used for measurements, including 64 EEG channels and 3 accelerometer channels. The sampling rate was set at 500 Hz, and an Acticap slim cap was used with a 10-20 layout (Ref = FCz, GND = AFz). The maximum impedance level of the electrodes was 5 kOhm, with impedance checks conducted both before and after measurements.

Several event markers were defined for the EEG recordings. The synchronization marker was labeled as “Sync,” and the start/stop trigger for EMG was represented by “S1.” Movement-related markers included the start of flexion (“S 64”), the start of extension (“S 32”), standard events (mid of movement, no error introduced, “S 48”), error introduction (“S 96”), and button press response (“S 80”). The experimental setup provided a controlled environment to study ERPs related to error detection. By using an active orthosis device and introducing deliberate errors, the dataset captured valuable insights into the underlying cognitive processes. The defined instructions for EEG measurements ensured precise recording and consistent representation of events, contributing to the reliability of the data and the robust foundation for the study’s objectives.

### EEG signal processing

EEG signal processing is essential in brain-computer interfaces (BCIs) and error-related potential studies (Gramann et al., 2014). Preprocessing employs a bandpass filter on the raw EEG signals, with a frequency range of 0.1Hz to 30Hz. The lower limit of 0.1Hz helps to attenuate slow drifts and artifacts, while the upper limit of 30Hz captures the essential frequency components related to cognitive activities. Bandpass filtering is vital in EEG signal processing as it reduces noise effects, focusing on the frequencies where the relevant information resides. This enhancement in the signal-to-noise ratio (SNR) preserves the information connected to the ERPs, facilitating a more accurate analysis.

Feature extraction transforms the preprocessed signals into a format that encapsulates the essential information. The following techniques were utilized: (1) Xdawn(Rivet et al., 2009) is a spatial filtering method that optimizes the SNR for ERPs, particularly in error detection. It enhances the discriminability between error and non-error states, offering robust feature representation. (2) ERPs are neurophysiological responses that reflect cognitive processes like error recognition(Kim et al., 2023). **Figure 1** presents the visualization for the ERPs in the 10-20 systems in our project. The P300 component, a positive deflection occurring approximately 300 milliseconds post-event, is a key feature in error detection. By extracting these specific ERP features, the analysis delves into underlying cognitive processes, contributing to the precise identification of error-related activities.

**Figure 1.**
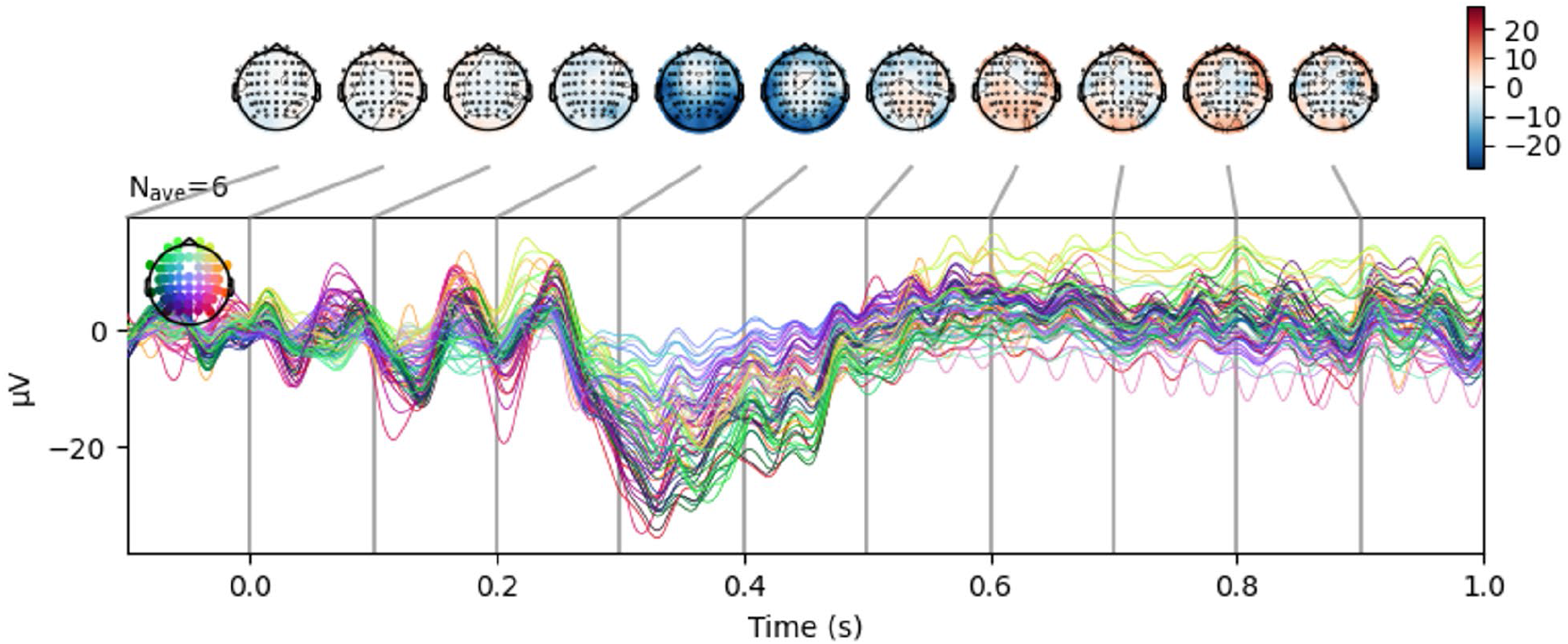
The example of ERPs and data visualization correspond to the average S 96 period of the 10-20 64 channels EEG system.

### Machine learning models

Our analysis employs two machine learning models: a customized ResNet and an ensemble model. We first create the training dataset by slicing out 1s trials of events (64 x 501) from the raw EEG sets. Both models take a 1s EEG trial (64 x 501) as input and predict it as either Error (S96) or Non-error (non-S96). A 0.1-30 Hz bandpass filter is applied to EEG raw data using the MNE package(*Tutorials — MNE 1.4.2 Documentation*, n.d.).

#### Customized ResNet model

The ResNet model starts with a 1-D convolutional layer, followed by batch normalization and a ReLU activation. A Residual Block structure is defined, which consists of two convolutional layers with batch normalization and ReLU activation, as well as a skip connection path from the input to the output. The residual block is applied once. After passing through the residual blocks, the output is passed to an adaptive average pooling layer, flattened, and finally fed to a fully connected linear layer for binary classification. **Figure 2** shows the architecture for the ResNet model. For training, the data is split into training, validation, and testing sets in a 7:1:2 ratio. The data is first preprocessed with a bandpass filter and subsequently standardized per channel across the entire training data. The model with the best validation loss during training is saved. Its performance is measured on the test set and its confusion matrix is plotted. The model is additionally evaluated using 10-fold cross-validation for each subject.

**Figure 2.**
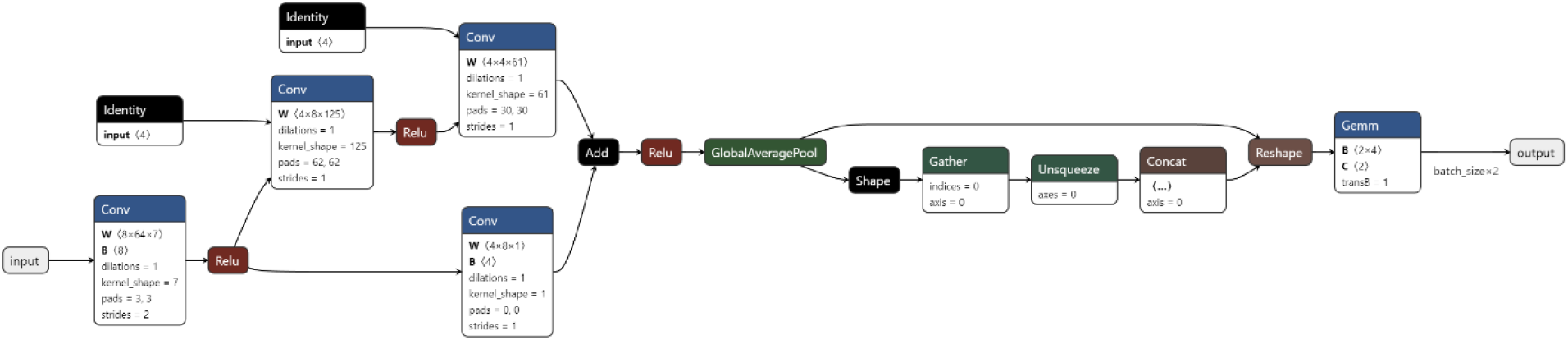
The architecture for the ResNet model. Conv block presents the 1D Convolution Layer and Gemm shows the Fully Connected Layer. The model has 8610 trainable parameters.

#### Ensemble model

An ensemble of machine learning models comprising MLP, logistic regression, SVM, random forest, and XGBoost are bundled in a pipeline with feature extraction and transformation steps. First, Xdawn Covariances(Barachant, n.d.) is applied to estimate the covariance matrices from the EEG data for each trial. This step applies the xDAWN spatial filtering method which enhances the signal-to-noise ratio, making it particularly suitable for ERP analysis. These covariance matrices are then transformed using Tangent Space. **Figure 3** shows the architecture for the Ensemble model. Because the covariance matrices lie in the space of Symmetric Positive Definite (SPD) matrices, which are not Euclidean, they are mapped to a tangent space where classical vector space operations can be performed. This allows the application of traditional machine learning techniques on the transformed data. The classifiers are combined into a soft-voting classifier, leveraging the averaged predicted probabilities for final decision-making. This ensemble model is evaluated using 10-fold cross-validation with the SMOTE (Synthetic Minority Over-sampling Technique) technique(Chawla et al., 2002) employed for each fold to oversample the minority class and address the class imbalance. Metrics, including accuracy, F1 score, and a confusion matrix, are calculated for each fold, visualized, and stored as text files, with the trained models saved as joblib files.

**Figure 3.**
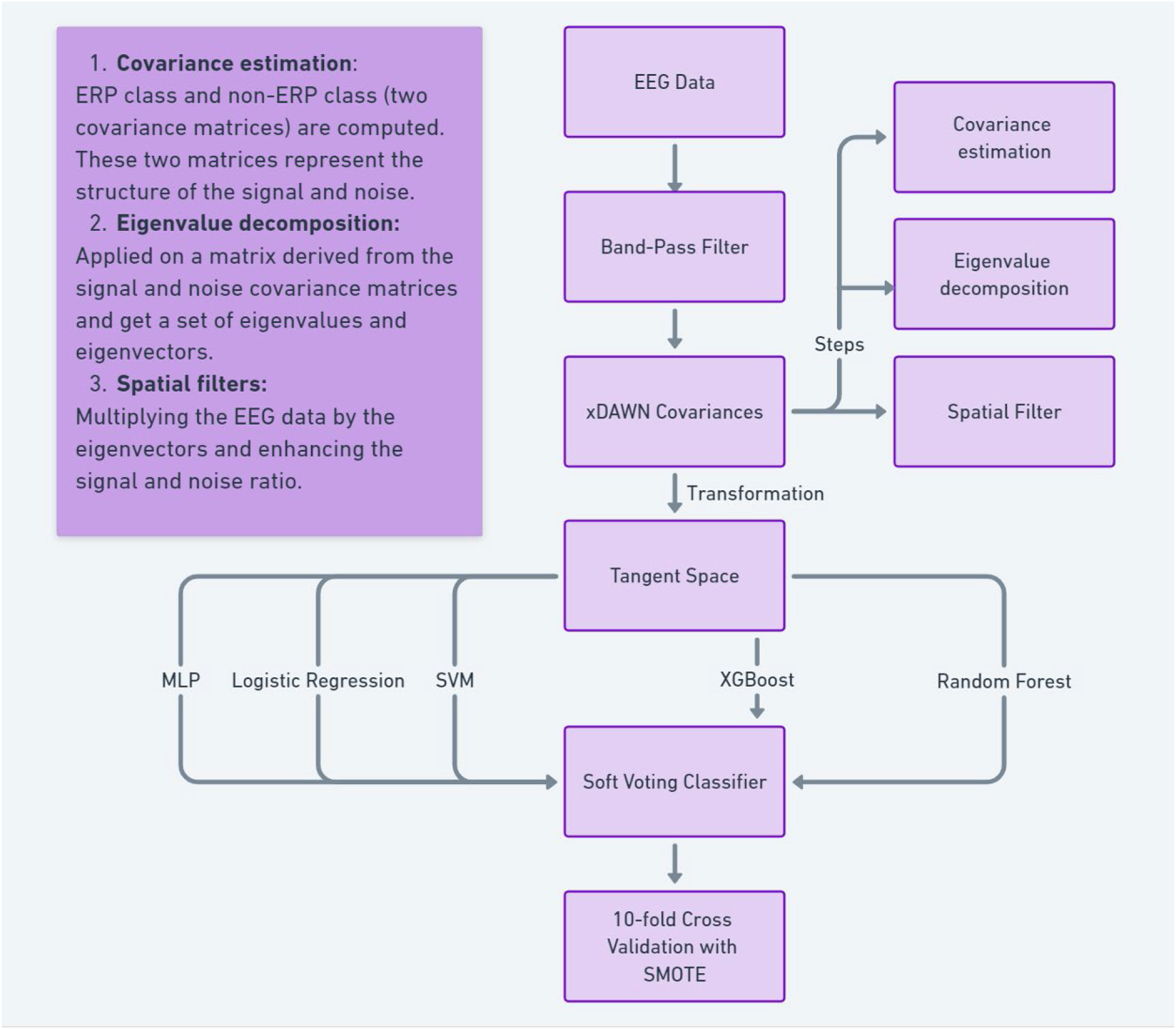
The architecture for the ensemble model. MLP is the Multilayer perceptron model. SVM is the Support Vector Machine model and XGBoost is the robust machine-learning algorithm.

#### Probability Map

The granularity of the probability map maintains a resolution of 0.1 seconds, and it is designed to forecast the likelihood of one-second intervals being classified either as S96 or non-S96 at the onset of every 0.1-second period. The algorithm achieves this by predicting the probabilities of the forthcoming one-second interval receiving either the S96 or non-S96 label and subsequently summing them. Within each 0.1-second interval, the pre-trained model computes the likelihood of the ensuing one-second period being assigned the S96 or non-S96 label.

## RESULTS

Ensemble and ResNet model were employed to detect erroneous behaviors during HRI. The results of both models are presented in terms of mean and standard deviation, total score, set score, and the number of true positives.

### Performance of Ensemble Model

**Table 1** shows the mean and standard deviation of the Ensemble model for different subjects. The standard deviation values are relatively small, indicating a consistent performance across the subjects. This consistency can be attributed to the soft-voting classifier, which neutralizes the individual results of the classifiers within the ensemble. The sum results in **Table 2** further highlight the efficiency of the Ensemble model, with a total score of 53804 and an online speed of 1.98 ms/data.

**Table 1.**
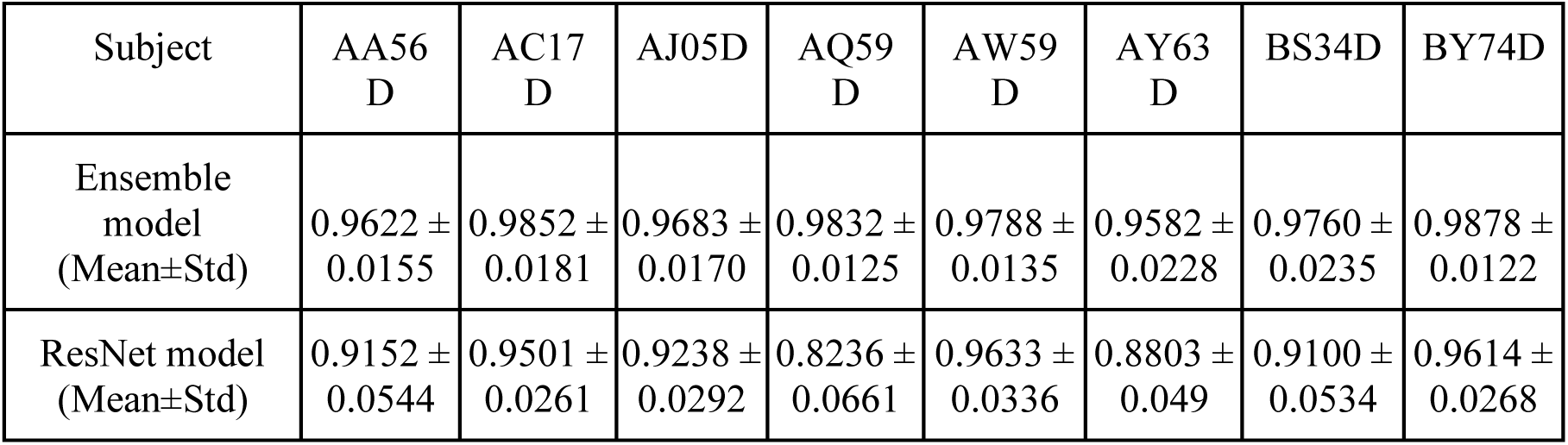
The results of 10-fold validation for the Ensemble model and ResNet model (Accuracy)

**Table 2.**
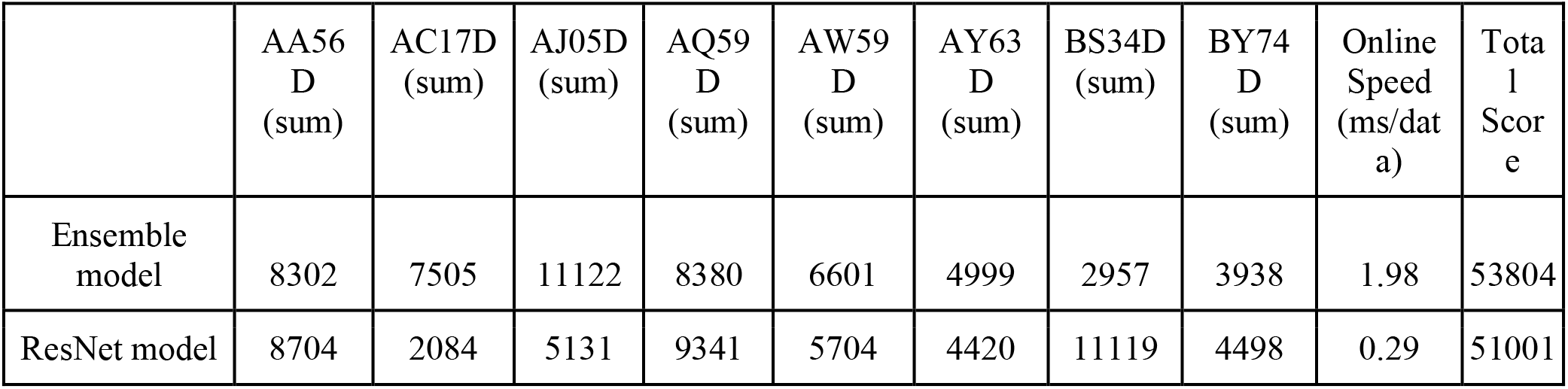
Test dataset results of Ensemble model and ResNet model.

|  | AA56<br>D<br>(sum) | AC17D<br>(sum) | AJ05D<br>(sum) | AQ59<br>D<br>(sum) | AW59<br>D<br>(sum) | AY63<br>D<br>(sum) | BS34D<br>(sum) | BY74<br>D<br>(sum) | Online<br>Speed<br>(ms/dat<br>a) | Tota<br>l<br>Scor<br>e |
| --- | --- | --- | --- | --- | --- | --- | --- | --- | --- | --- |
| Ensemble<br>model | 8302 | 7505 | 11122 | 8380 | 6601 | 4999 | 2957 | 3938 | 1.98 | 53804 |
| ResNet model | 8704 | 2084 | 5131 | 9341 | 5704 | 4420 | 11119 | 4498 | 0.29 | 51001 |

### Performance of ResNet Model

The ResNet model’s mean and standard deviation values in **Table 1** reveal a performance that varies across subjects. Although the model achieved commendable results, there is a noticeable disparity in some cases, signifying room for potential improvement. The sum results for the ResNet model, presented in **Table 2**, yield a total score of 51001, with a significantly faster online speed of 0.29 ms/data compared to the Ensemble model.

### Comparison of Ensemble and ResNet Models

**Figure 4** presents the confusion matrix by using all of the data. Comparing the results of the Ensemble and ResNet models, the Ensemble model generally outperforms the ResNet model in terms of mean values and consistency. However, the ResNet model excels in online processing speed. **Figure 5** presents the example error time point prediction using the probability map. This calculated probability is then allocated to the imminent ten data points, each symbolizing a fraction of one-tenth of a second. The algorithm perpetually revises the maximum probability for each point throughout this process. In the concluding phase, the algorithm pinpoints the six time points that generate the highest probabilities.

**Figure 4.**
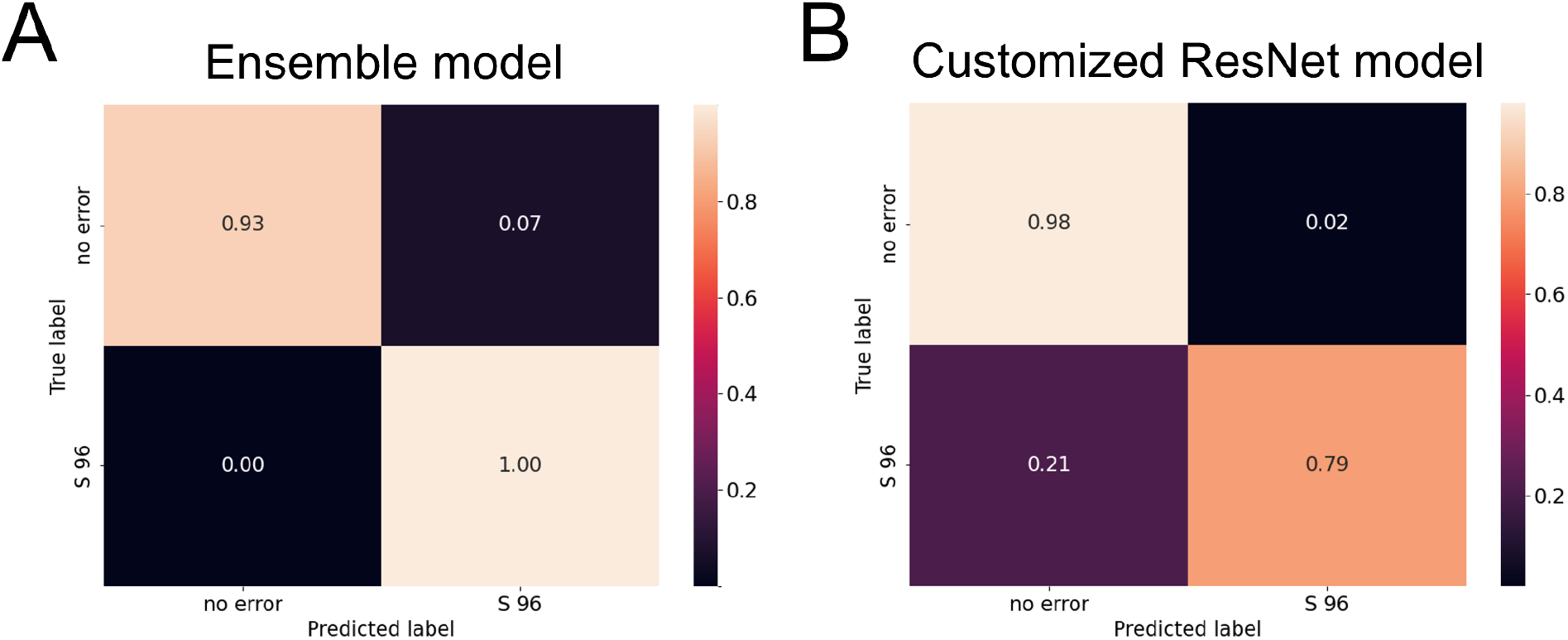
The result of the confusion matrix using all data, A is the result of the confusion matrix by using the Ensemble model, and B is the result of the confusion matrix by using the ResNet model.

**Figure 5.**
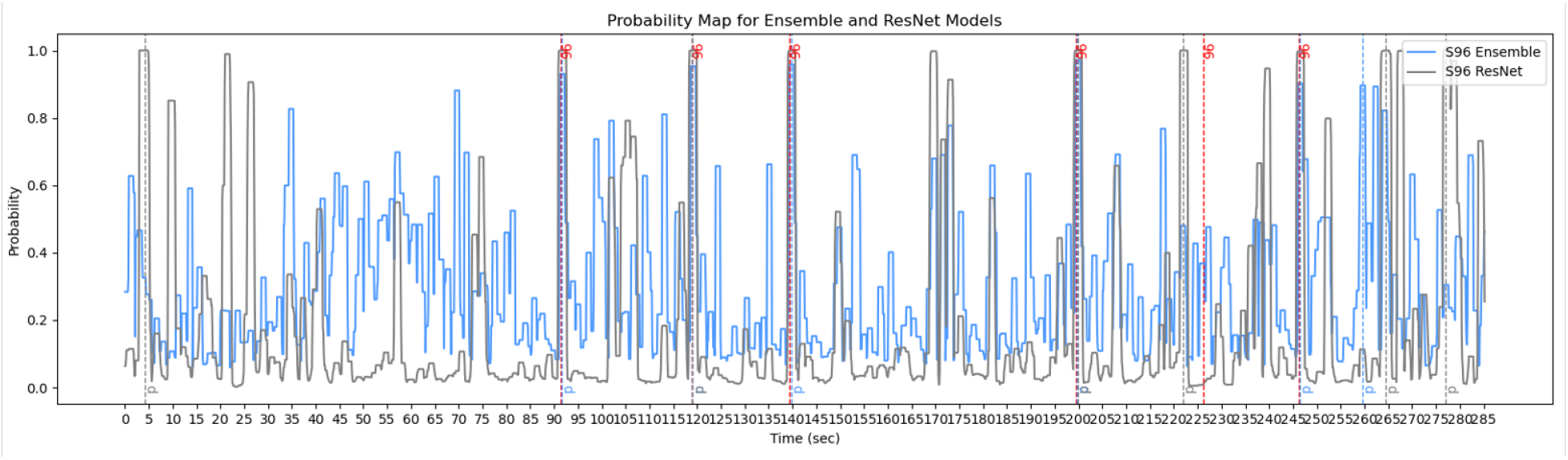
The example error time point prediction using the probability map, the red dotted line indicates ground truth error time, the blue line indicates the prediction for the Ensemble model, and the black line indicates the prediction for the ResNet model.

### Offline Stage Evaluation

The evaluation of the offline stage competition was conducted based on the comparison of the submitted sample indices against the ground truth. Each of the submitted sample indices (six per set) was converted into corresponding time points in milliseconds and then compared against the reference. The temporal error, defined as the difference in milliseconds, was calculated up to a maximum time point limit (e.g., 3000 ms). If a detection occurred outside this maximum time point, a penalty equivalent to the maximum time point limit was added to the team’s total. The error term was summed across all sets and subjects, resulting in a grand total that determined the winners of the competition. This rigorous evaluation process ensured an accurate and fair assessment of the models’ capabilities in the offline stage, reflecting their efficiency and effectiveness in detecting error-related potentials.

### Offline Stage Results

**Tables 3** and **4** provide detailed results of the offline stage, including predicted labels, ground truth labels, set scores, and the number of true positives. **Figure 6** presents the result of the offline stage competition where our method secured first place. The Ensemble model achieved a sum score of 53804 with 48 true positives, while the ResNet model achieved a sum score of 51001 with 52 true positives. It is important to note that the score mentioned here is different from the one presented in **Table 3**. The discrepancy arises from improvements made to the Ensemble model. Specifically, we refined the model by deleting some of the worse-performing individual classifiers within the ensemble, leading to enhanced accuracy and a lower score, indicating better performance.

**Figure 6.**
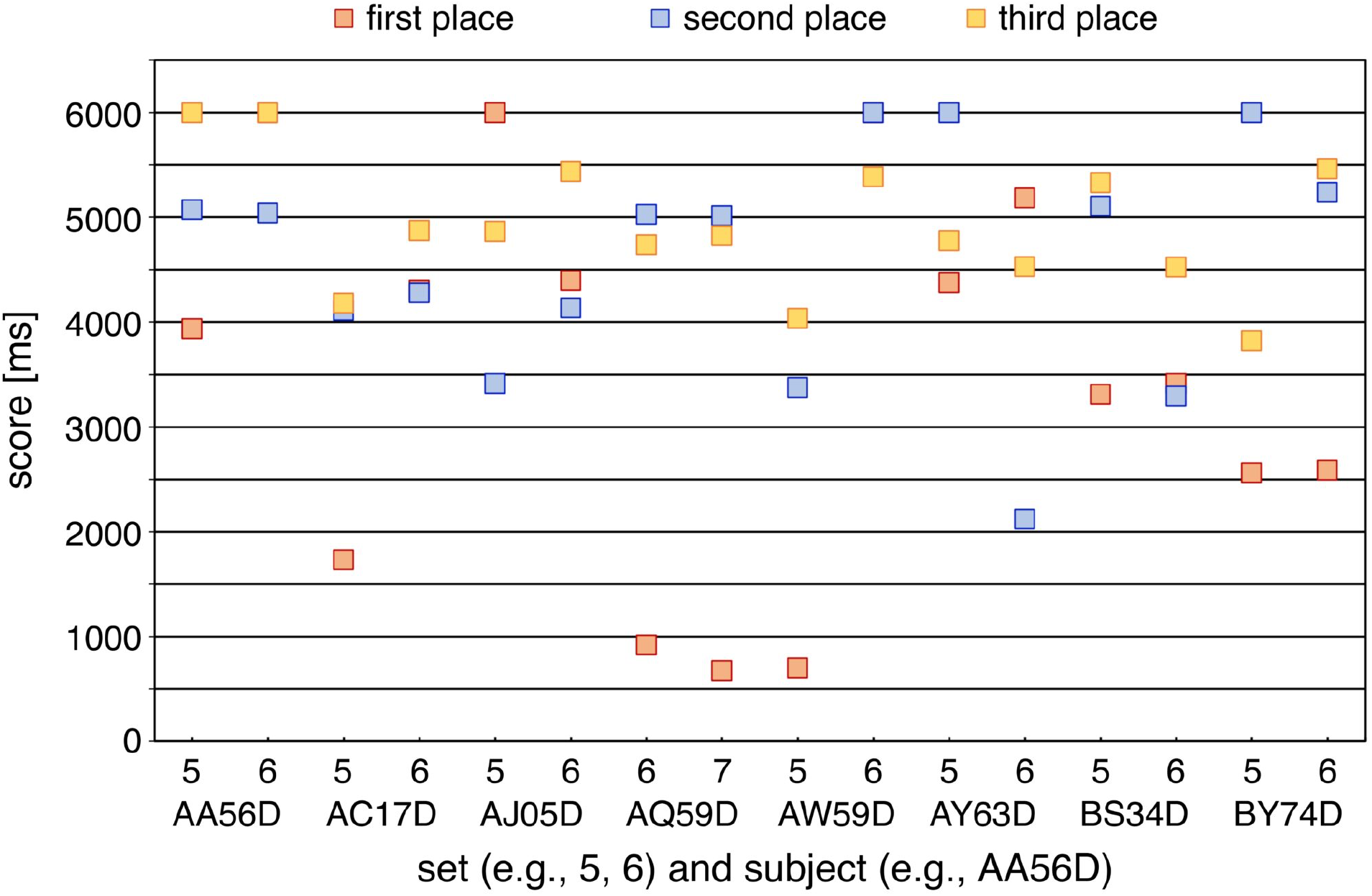
The result of the offline stage competition showcased our data’s achievement in the first place. The figure illustrates the performance rankings and highlights our superior performance in this challenging domain.

**Table 3.** The results of the offline stage include the predicted label and ground truth label for the Ensemble model.

| DataSet | Predicted | Ground Truth | Set Score | No. of True Positives |
| --- | --- | --- | --- | --- |
| AQ59D_set6 | 45470, 75920, 83870, 95020, 105870, 116620 | 30989, 45383, 80984, 94935, 105786, 116483 | 3306 | 3 |
| AQ59D_set7 | 32444, 51294, 62494, 73394, 114194, 119544 | 20550, 32360, 51188, 62390, 73316, 119470 | 5074 | 1 |
| AJ05D_set5 | 21290, 25740, 51190, 59490, 73540, 94490 | 29859, 44867, 59376, 73436, 94344, 111985 | 6000 | 0 |
| AJ05D_set6 | 34594, 58044, 98844, 111294, 116794, 130544 | 34472, 46150, 57921, 98744, 111208, 130666 | 5122 | 1 |
| BS34D_set6 | 39644, 56694, 66894, 72494, 93594, 113694 | 39528, 56616, 66819, 83440, 93522, 113575 | 1460 | 5 |
| BS34D_set5 | 29868, 40068, 50368, 84018, 89218, 110918 | 29753, 39928, 50250, 83927, 94285, 110885 | 1497 | 5 |
| AW59D_set5 | 28425, 53625, 72825, 82425, 95075, 104875 | 28346, 53529, 72696, 82370, 94985, 104723 | 601 | 6 |
| AW59D_set6 | 89920, 115820, 124870, 151470, 156720, 169170 | 80025, 89766, 99353, 115699, 126243, 151322 | 6000 | 0 |
| AA56D_set5 | 52545, 80595, 94095, 106845, 110545, 123245 | 38339, 52386, 80472, 93949, 110389, 127213 | 5156 | 1 |
| AA56D_set6 | 41745, 46345, 56245,<br>76595, 108395, 112445 | 35438, 45679, 56132,<br>76437, 92538, 112236 | 3146 | 4 |
| BY74D_set5 | 35958, 45858, 62708,<br>71708, 98358, 107908 | 35875, 45760, 55458,<br>71575, 84799, 107807 | 2415 | 4 |
| BY74D_set6 | 63259, 79809, 99809,<br>116559, 126709, 147159 | 63161, 79681, 99674,<br>116457, 126649, 143375 | 1523 | 5 |
| AY63D_set5 | 58813, 76263, 92213,<br>100713, 104713, 116163 | 30827, 44573, 58691,<br>76108, 104608, 116030 | 4238 | 2 |
| AY63D_set6 | 33947, 48347, 68697,<br>88947, 106247, 123597 | 33830, 48181, 68574,<br>88862, 106121, 123453 | 761 | 6 |
| AC17D_set6 | 42913, 59413, 86063,<br>99213, 106263, 116063 | 32985, 42838, 59319,<br>85930, 99094, 115967 | 5096 | 1 |
| AC17D_set5 | 44941, 49341, 65591,<br>75941, 85941, 106041 | 24574, 44832, 65498,<br>75827, 85830, 105950 | 2409 | 4 |
| Sum |  |  | 53804 | 48 |

**Table 4.** The result of the offline stage includes the predicted label and ground truth label for the ResNet model.

| DataSet | Predicted | Ground Truth | Set Score | No. of True Positives |
| --- | --- | --- | --- | --- |
| AQ59D_set6 | 31120, 36170, 81120,<br>94270, 110720, 115220 | 30989, 45383, 80984,<br>94935, 105786, 116483 | 4267 | 2 |
| AQ59D_set7 | 32594, 53794, 73344,<br>89494, 100894, 119544 | 20550, 32360, 51188,<br>62390, 73316, 119470 | 5074 | 1 |
| AJ05D_set5 | 29990, 53490, 59590,<br>73540, 94540, 118540 | 29859, 44867, 59376,<br>73436, 94344, 111985 | 2645 | 4 |
| AJ05D_set6 | 34544, 58044, 91044,<br>98894, 111294, 130844 | 34472, 46150, 57921,<br>98744, 111208, 130666 | 2486 | 4 |
| BS34D_set6 | 54294, 67044, 83494,<br>93594, 108394, 113694 | 39528, 56616, 66819,<br>83440, 93522, 113575 | 5119 | 1 |
| BS34D_set5 | 16668, 29868, 85318,<br>94518, 106668, 110818 | 29753, 39928, 50250,<br>83927, 94285, 110885 | 6000 | 0 |
| AW59D_set5 | 28375, 53625, 72875,<br>82375, 92725, 104875 | 28346, 53529, 72696,<br>82370, 94985, 104723 | 1461 | 5 |
| AW59D_set6 | 80170, 87570, 115770,<br>126420, 142070, 151420 | 80025, 89766, 99353,<br>115699, 126243, 151322 | 4243 | 2 |
| AA56D_set5 | 38495, 52545, 94095,<br>110595, 123245, 127345 | 38339, 52386, 80472,<br>93949, 110389, 127213 | 3447 | 3 |
| AA56D_set6 | 35695, 56295, 62845,<br>72045, 76595, 110995 | 35438, 45679, 56132,<br>76437, 92538, 112236 | 5257 | 1 |
| BY74D_set5 | 35958, 43658, 53808,<br>72158, 84958, 107908 | 35875, 45760, 55458,<br>71575, 84799, 107807 | 2926 | 4 |
| BY74D_set6 | 63309, 79809, 109159,<br>116609, 126659, 143509 | 63161, 79681, 99674,<br>116457, 126649, 143375 | 1572 | 5 |
| AY63D_set5 | 30963, 44813, 59913,<br>104663, 110913, 116163 | 30827, 44573, 58691,<br>76108, 104608, 116030 | 3509 | 3 |
| AY63D_set6 | 33947, 48347, 68697,<br>89097, 106247, 123597 | 33830, 48181, 68574,<br>88862, 106121, 123453 | 911 | 6 |
| AC17D_set6 | 33113, 42913, 59413,<br>86063, 99313, 112413 | 32985, 42838, 59319,<br>85930, 99094, 115967 | 1649 | 5 |
| AC17D_set5 | 24641, 44891, 65591,<br>75891, 85891, 106041 | 24574, 44832, 65498,<br>75827, 85830, 105950 | 435 | 6 |
| Sum |  |  | 51001 | 52 |

The final standings in the offline stage competition are summarized in **Table 5**, which shows the accumulated scores in milliseconds for the top three teams. Analysis of the competition results reveals that our team secured first place with an accumulated score of 56132 ms, outperforming the second and third-place teams by significant margins. The difference in scores between the top three teams highlights the efficiency and accuracy of our detection models, particularly in the context of time-constrained error detection. The success in the competition underscores the effectiveness of our approach, validates the methodologies employed, and represents a significant milestone in our research into Human-Robot Interaction (HRI).

**Table 5.** The result of the offline stage competition for the top three teams.

|  | 1st place team | 2nd place team | 3rd place team |
| --- | --- | --- | --- |
| Accumulated Score<br>(ms) | 56132 | 73270 | 78820 |

## DISCUSSION

In this research, we have studied and compared the Ensemble and ResNet models for detecting erroneous behaviors during Human-Robot Interaction (HRI). Our findings corroborate with previous research(Barachant, n.d.; Craik et al., 2019) that Ensemble models, through the use of soft-voting classifiers, can provide consistent performance across different subjects, making them suitable for error detection in HRI. However, the ResNet model’s superior online processing speed poses an interesting trade-off between accuracy and real-time responsiveness. This insight aligns with the growing need for efficient, real-time error detection in adaptive and intuitive robotic systems. The offline stage competition further validates our approach and demonstrates the practical applicability of our models. The competition’s evaluation methodology, involving the conversion of sample indices to corresponding time points and calculation of temporal error, has provided a rigorous assessment of our models’ performance. We’ve managed to improve the Ensemble model’s performance by deleting some of the worst-performing models, leading to a lower score and better performance. Future work may include further optimization of these models and exploration of additional ensemble techniques to enhance the detection of error-related potentials.

Deep learning has significantly revolutionized the way machine learning is perceived in the HRI domain(Fitz & Romero, 2021; Karoly et al., 2021). Many previous works utilize neural networks for EEG signal classification, with the majority of the networks CNN-based(Craik et al., 2019). The primary design features for CNNs are the number of convolutional layers and the type of end classifier, which is typically a fully connected layer. For example, Lun et al. (Lun et al., 2020) designed a 5-layer CNN with max-pooling and a fully connected layer for a motor imagination classification task. Their network is able to obtain State-of-the-art performance on the Physionet database with 97.28 accuracy. ResNet(He et al., 2016) is a variant of CNN that has been the state-of-the-art model for image classification in computer vision. By introducing skip connections, ResNet makes error propagation during training easier. In our work, following the path of existing works, we designed a lightweight ResNet model for EEG classification. Another line of work involves expert feature extraction methods for CNN classification. Continuous wavelet transforms(Mao et al., 2020), Power spectral density(R. Wang et al., 2015), etc. have been used to extract features for EEG classification. xDawn(Rivet et al., 2009) is a state-of-the-art feature extractor that is specifically designed for processing ERPs in EEG data. It aims to maximize the signal-to-noise ratio of the ERP components thus enhancing their detectability. We used the xDawn algorithm to enhance ERP detectability and used an ensemble of simple classifiers to do the classification.

## CONCLUSION

This study provides valuable insights into the detection of error-related potentials in Human-Robot Interaction, with a focus on Ensemble and ResNet models. Our models’ performance in various sets and subjects has been rigorously evaluated, and the offline stage competition attests to the robustness and reliability of our approach. Our research contributes to ongoing efforts in the development of adaptive and intuitive robotic systems, laying the groundwork for more sophisticated, real-time interactions between humans and robots.

## DATA AND CODE AVAILABILITY STATEMENT

All the datasets used in this study are available at https://zenodo.org/record/7951044 for training data, and at https://zenodo.org/record/7966275 for testing data. All the source codes of models are available at https://github.com/NeuroPrior-AI/IntEr-HRI-Competition.

## AUTHOR CONTRIBUTIONS

ZR: methodology, software, formal analysis, writing—original draft, and visualization. XX: methodology, software, formal analysis, writing—original draft, and visualization. YT: methodology, software, formal analysis, writing—original draft, and visualization. BZ: conceptualization, methodology, software. CPW: investigation, visualization. DX: conceptualization, methodology, resources, investigation, supervision, writing—original draft, and visualization. All authors contributed to the article and approved the submitted version.

## ACKNOWLEDGMENTS

We would like to thank all the members of the NeuroPrior AI, Neurointelligence Labs, whose collaboration and dedication have contributed immensely to the success of this work. We are grateful to the organizers of the International Joint Conference on Artificial Intelligence (IJCAI) Human-Robot Interaction (HRI) Competition for providing the platform to showcase our research. The authors would also like to acknowledge the support from various funding agencies, colleagues, and anyone else who contributed to the project.

